# Benefits of improved air quality on aging lungs – Impacts of genetics and obesity

**DOI:** 10.1101/521328

**Authors:** Anke Hüls, Dorothea Sugiri, Michael J Abramson, Barbara Hoffmann, Holger Schwender, Katja Ickstadt, Ursula Krämer, Tamara Schikowski

## Abstract

**Introduction:** The beneficial effect of improving air quality on lung function in the elderly remains unclear. We examined associations between decline in air pollutants and lung function and effect modifications by genetics and BMI in elderly German women.

**Methods:** Data were analysed from the prospective SALIA cohort study (N=601). Spirometry was conducted at baseline (1985-1994; 55 years), in 2007-2010 and in 2012-2013. Air pollution concentrations at home addresses were determined for each time point using land-use regression models. GLI z-scores were calculated. Weighted genetic risk scores (GRS) were determined from lung function-related risk alleles and used to investigate interactions with improved air quality. Adjusted linear mixed models were fitted.

**Results:** Air pollution levels decreased substantially during the study period. Reduction of air pollution was associated with an increase of z-scores for FEV_1_ and FEV_1_/FVC. For a decrease of 10 μg/m^3^ in NO_2_, the z-score for FEV_1_ increased by 0.14 (95%CI: 0.01; 0.26). However, with an increasing number of lung function-related risk alleles, the benefit from improved air quality decreased (GRSxNO_2_-interaction:p=0.029). Interactions with BMI were not significant.

**Conclusions:** Reduction of air pollution is associated with a relative improvement of lung function in elderly women, but also depends on their genetic make-up.

## Introduction

Air pollution is a major environmental risk factor affecting a range of acute and chronic aging-related conditions such as pulmonary and cardiovascular diseases worldwide [1, 2]. Previous studies showed that long-term exposure to air pollution has adverse effects on chronic respiratory diseases [3–5] as well as lung function [3, 6–11].

However, the beneficial effects of improved air quality on the respiratory system are less well understood. Studies from Switzerland and Germany provided evidence that improvement of air quality could have a beneficial effect on respiratory symptoms [12, 13]. In addition, improved air quality in southern California was shown to be associated with improvements in childhood lung-function growth [14] and improved air quality in Switzerland with an attenuated decline in lung function among adults [15]. However, most research has been conducted by including data from only two time points of investigation and little is known about the beneficial effect of a reduction in air pollutants on aging lungs of elderly people.

Beside air pollution, studies provide evidence that the genetic make-up [16] as well as abdominal obesity [17] play a major role for impaired lung function. However, little is known about the underlying mechanisms and about the impacts of genetics and obesity on the beneficial effects of improved air quality on aging lungs. Thun et al. [18] performed a gene-environment (GxE) interaction analysis in the Swiss study on Air Pollution And Lung Disease in Adults (SAPALDIA) to investigate the genetic make-up associated with attenuated lung function decline due to improvement of air quality. However, after correction for multiple testing none of the lung function-associated single nucleotide polymorphisms (SNPs) they included in these analyses modified the association between improvement of air quality and attenuated lung function decline. In the same cohort, Schikowski et al. [19] showed that improved air quality was associated with attenuated age-related reductions in lung function over time among low- and normal-BMI participants, but not in overweight or obese participants.

Improvements in air quality over time provide the backdrop for a “natural experiment” to examine the potential beneficial health effects. The SALIA cohort (***S***tudy on the influence of ***A***ir pollution on ***L***ung function, ***I***nflammation and ***A***ging) offers a well-characterized exposure history of the study participants with an on-going improvement of air quality over time, as well as consistently assessed health data and lung function of the study participants at up to three time points. This cohort study was initiated 28 years ago and the participating women were up to 83 years old at the last follow-up investigation.

We examined the association between improved air quality and age-related lung function decline in elderly women from the SALIA cohort at three time points of examination (1985-1994, 2007-2010, and 2012-2013) and effect modifications by genetics and BMI.

## Methods

More details on the methods are given in the supplementary material.

### Study design and study population

The SALIA study population was a sample of women from the urban Ruhr area and the adjacent rural Muensterland, Germany. Between 1985 and 1994, health examinations and lung function measurements were conducted in 2785 women [20].

The first follow-up examination was conducted between 2007 and 2010 [21] and the second between 2012 and 2013 [22]. In total, 834 women participated in the first follow-up examination and 542 in the second. Ethical approval was obtained from the Ethical Committees of the University of Bochum and the Heinrich Heine University. We received written informed consent from all participants.

### Assessment of air pollution

Outdoor air pollution (NO_2_, NO_x_, PM_2.5_ and PM_10_) concentrations were assessed according to the ESCAPE (European Study of Cohorts for Air Pollution Effects) protocol [23, 24]. Air pollution was monitored over one year (2008-2009) in the study area. Land-use regression (LUR) models were applied to the home addresses.

To characterize the levels of air pollution exposure at each time point of investigation, we used extrapolation procedures. Ratios were calculated between air pollution concentrations at government background monitoring stations during a two-year period around each time point of investigation and the period of ESCAPE monitoring campaign. The values from the LUR models were corrected for spatial trends by multiplying them with these ratios. The implicit assumption of proportional spatial contrasts over time was validated with data from six routine monitoring stations situated in the study area and covering the investigation period.

### Assessment of genotypes

A genome-wide genotyping was performed in 462 women who participated in the follow-up investigation 2007-2010, using the Affymetrix Axiom™ Precision Medicine Research Array. SNPs were imputed on the 1000 Genomes reference panel (Phase III) using Minimac3 [25].

### Assessment of pulmonary function

Spirometry was performed according to the ATS/ETS recommendations [26] at all three time points. Forced expiratory volume in one second (FEV_1_) and forced vital capacity (FVC) were measured. To control for age and height-dependency of lung function, we calculated z-scores from the Global Lung Initiative (GLI) reference values [27]. In a previous publication, we showed that GLI z-scores fitted cross-sectionally and longitudinally with FEV_1_, FVC and FEV_1_/FVC measured in our SALIA cohort [22].

### Statistical analysis

We investigated the association between long-term improvement of air quality between baseline and first follow-up and between baseline and second follow-up with the corresponding change in GLI z-scores for FEV_1_, FVC and FEV_1_/FVC in linear mixed models with random participant intercepts [28]. *A priori* selected covariates at three time points which could potentially act as confounders included age, BMI at baseline, change of BMI, highest educational status of participant or husband, smoking status and exposure to second hand smoke (SHS).

To investigate the impact of the genetic make-up on the association between improvement of air quality with change in lung function, we conducted a gene-environment interaction analysis including 49 SNPs shown to be associated with impaired lung function in genome-wide association studies (GWAS) [16]. To summarize the genetic risk factors for impaired lung function, we used weighted genetic risk scores (GRS), which aggregate measured genetic effects and therefore increase the power to detect gene-environment interactions [29, 30]. External weights (β-estimates for marginal genetic effects) were gained from the GWAS with largest sample size available [16]. GRS were calculated for each individual by multiplying the number of risk alleles for each of the 49 SNPs with the respective external weights and calculating the sum over all SNPs [30]. We estimated the interaction of this GRS with improved air quality in adjusted linear mixed models with a random participant intercept. Additionally, associations between genotype status at the lung function related single SNPs (additive model) and change in lung function z-scores were estimated adjusted for principal components (PCs) 1–10 to adjust for possible population stratification and age as covariates.

Finally, we investigated the interaction between the participants’ average BMI from baseline to second follow-up and improvement of air quality on change of lung function. Associations between improvement of air quality and change in lung function were stratified by tertiles of average BMI.

All statistical analyses were carried out with R 3.4.3 for Windows [31].

## Results

### Description of study participants, air pollution and pulmonary function

The characteristics of the study participants are described in Table 1. All women with lung function measurements at baseline and at least at one follow-up investigation (2007-2010 and/or 2012-2013) were included in the analysis (N=601). Of these, we had data from 587 women at first follow-up and 333 at second follow-up.

**Table 1:** Description of study population at baseline, first and second follow-up (including lung function). All women with lung function measurements at ≥2 time points of measurement were included in the analysis (N=601).

|  | **Baseline**<br>**(1985-1994)** | **1<sup>st</sup> Follow-up**<br>**(2007-2010)** | **2<sup>nd</sup> Follow-up**<br>**(2012-2013)** |
| --- | --- | --- | --- |
| N | 601 | 587 | 333 |
| Age years, mean ± sd | 54.3 ± 0.8 | 73.3 ± 3.3 | 77.4 ± 3.4 |
| BMI kg/m<sup>2</sup>, mean ± sd | 26.7 ± 3.9 | 27.3 ± 4.5 | 28.5 ± 4.5 |
| Change from baseline BMI*, mean ± sd | | 0.7 ± 2.7 | 2.0 ± 3.0 |
| >10 years of education, n (%) | 203 (33.8%) | 200 (34.1%) | 117 (35.1%) |
| 10 years of education, n (%) | 285 (47.4%) | 277 (47.2%) | 148 (44.4%) |
| <10 years of education, n (%) | 111 (18.5%) | 108 (18.4%) | 68 (20.4%) |
| Smoker, n (%) | 62 (10.3%) | 19 (3.2%) | 13 (3.9%) |
| Ex-smoker, n (%) | 61 (10.1%) | 99 (16.9%) | 48 (14.4%) |
| Never smoker, n (%) | 477 (79.4%) | 469 (79.9%) | 272 (81.7%) |
| Passive smoking, n (%) | 276 (45.9%) | 350 (59.6%) | 205 (61.6%) |
| GLI z-score FEV<sub>1</sub>, mean ± sd | -0.2 ± 1.0 | 0.2 ± 1.0 | 0.1 ± 1.1 |
| Change from baseline GLI z-score FEV<sub>1</sub><sup>†</sup>, mean ± sd | | 0.4 ± 0.8 | 0.3 ± 0.8 |
| GLI z-score FVC, mean ± sd | 0.0 ± 0.9 | 0.3 ± 0.9 | 0.5 ± 1.0 |
| Change from baseline GLI z-score FVC<sup>†</sup>, mean ± sd | | 0.3 ± 0.7 | 0.4 ± 0.8 |
| GLI z-score FEV<sub>1</sub>/FVC, mean ± sd | -0.4 ± 0.8 | -0.2 ± 0.9 | -0.7 ± 0.8 |
| Change from baseline GLI z-score FEV<sub>1</sub>/FVC<sup>†</sup>, mean ± sd | | 0.2 ± 0.9 | -0.3 ± 0.8 |

\*Change of BMI between baseline and 1<sup>st</sup> follow-up (BMI at 1<sup>st</sup> follow-up - BMI at baseline) and between baseline and 2<sup>nd</sup> follow-up (BMI at 2<sup>nd</sup> follow-up - BMI at baseline).
<sup>†</sup>Change of lung function between baseline and 1<sup>st</sup> follow-up (lung function at 1<sup>st</sup> follow-up - lung function at baseline) and between baseline and 2<sup>nd</sup> follow-up (lung function at 2<sup>nd</sup> follow-up - lung function at baseline).

The average age of these women at baseline investigation in the years 1985-1994 was 54.3 years, 73.3 at first follow-up in 2007-2010, and 77.4 at second follow-up investigation in 2012-2013. Mean GLI z-scores for FEV_1_ and FVC increased from baseline to both follow-up investigations, whereas mean z-scores for FEV_1_/FVC only increased from baseline to first follow-up, but decreased from baseline to second follow-up. The same was observed in the analysis of complete cases with available lung function measurements at all three time points of measurement (Table S1).

The distributions of air pollutants during the study period are presented in Figure 1 and Table 2. Air pollution levels fell during the study period (e.g., NO_2_ from a median of 33.4 to 19.7 μg/m^3^). At both follow-up investigations, PM_2.5_ and PM_10_ levels were below the European Union (EU) limit values for a one-year averaging period, whereas NO_2_ levels still exceeded the EU limit values in some urban areas (Figure 1). Since we used the ratio with baseline PM_10_ measurements for the back-extrapolation of PM_10_ as well as PM_2.5_, reduction of PM_2.5_ was strongly correlated with reduction of PM_10_ (e.g. r^2^=0.98 between baseline and 1^st^ follow-up). Furthermore, reduction of NO_2_ was strongly correlated with reduction of NO_x_, because extrapolation of NO_x_ was based on the ratio of NO_2_ measurements (e.g. r^2^=0.88 between baseline and 1^st^ follow-up), whereas the correlation between reduction of PM and nitrogen oxides was moderate (e.g., r^2^=0.63 for reduction of PM_10_ and NO_x_ between baseline and 1^st^ follow-up) (Table S2).

**Figure 1:**
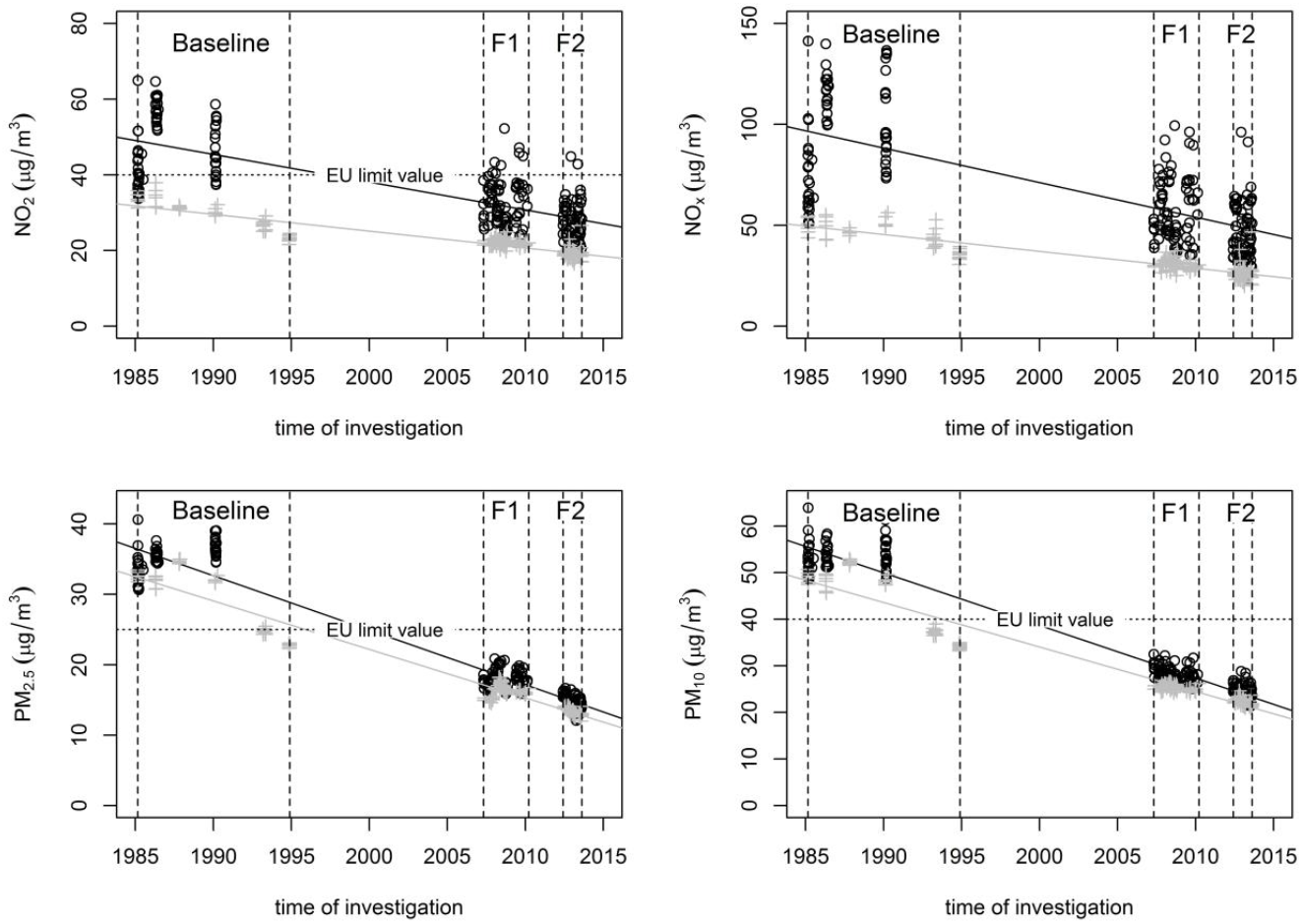
Improvement of air quality during the study period (1985-2013) Air quality standards: http://ec.europa.eu/environment/air/quality/standards.htm; European Union (EU) limit values for a one-year averaging period. No EU limit values available for NO_x_. Black: urban area, grey: rural area; F1: 1^st^ Follow-up investigation, F2: 2^nd^ Follow-up investigation All women with lung function measurements at ≥2 time points of measurement were included in the analysis (N=601).

**Table 2:** Description of air pollution exposures during the study period (1985-2013).

|  | **Baseline (1985-1994)** |  | **1<sup>st</sup> Follow-up (2007-2010)** |  | **2<sup>nd</sup> Follow-up (2012-2013)** |  |
| --- | --- | --- | --- | --- | --- | --- |
|  | **Median (IQR)** | **Min; Max** | **Median (IQR)** | **Min; Max** | **Median (IQR)** | **Min; Max** |
| NO<sub>2</sub> (µg/m<sup>3</sup>) | 33.4 (13.8) | 20.3; 84.1 | 23.4 (7.5) | 18.4; 68.2 | 19.7 (6.3) | 16.1; 46.1 |
| NO<sub>x</sub> (µg/m<sup>3</sup>) | 52.8 (39.0) | 26.8; 184.1 | 34.1 (19.7) | 21.7; 116.1 | 27.7 (16.1) | 18.7; 98.9 |
| PM<sub>2.5</sub> (µg/m<sup>3</sup>) | 33.0 (4.8) | 22.0; 41.3 | 16.6 (1.7) | 14.2; 21.4 | 13.9 (1.5) | 12.0; 17.9 |
| PM<sub>10</sub> (µg/m<sup>3</sup>) | 49.9 (8.0) | 32.2; 65.1 | 26.0 (2.2) | 23.2; 33.2 | 22.9 (1.8) | 20.3; 28.9 |
| Improvement of NO<sub>2</sub> from baseline* (µg/m<sup>3</sup>) |  |  | 9.4 (7.5) | -29.2; 41.3 | 12.6 (7.0) | -7.3; 44.2 |
| Improvement of NO<sub>x</sub> from baseline* (µg/m<sup>3</sup>) |  |  | 20.0 (18.3) | -39.6; 107.2 | 24.7 (21.8) | -19.6; 113.1 |
| Improvement of PM<sub>2.5</sub> from baseline* (µg/m<sup>3</sup>) |  |  | 16.4 (5.6) | 5.6; 23.6 | 19.2 (3.5) | 8.6; 28.6 |
| Improvement of PM<sub>10</sub> from baseline* (µg/m<sup>3</sup>) |  |  | 23.6 (8.8) | 6.9; 36.4 | 26.6 (5.0) | 9.8; 42.8 |

All women with lung function measurements at $\geq 2$ time points of measurement were included in the analysis (N=601).
\*Improvement of air quality between baseline and 1<sup>st</sup> follow-up (air pollution at baseline – air pollution at 1<sup>st</sup> follow-up) and between baseline and 2<sup>nd</sup> follow-up (air pollution at baseline – air pollution at 2<sup>nd</sup> follow-up).

### Improvement of air quality and change in lung function

Reduction of air pollution was associated with an increase of z-scores for FEV_1_ and FEV_1_/FVC, but not for FVC (Figure 2 and Table S3). A decrease of 10 μg/m^3^ in NO_2_ and 20 μg/m^3^ in NO_x_ over the study period was associated with an increase of FEV_1_ and FEV_1_/FVC (e.g. for NO_2_ and FEV_1_ by 0.14 (95%CI 0.01; 0.26)). For a decrease of 10 μg/m^3^ in PM_10_, the z-score for FEV_1_/FVC increased by 0.21 (95%CI 0.07; 0.35).

**Figure 2:**
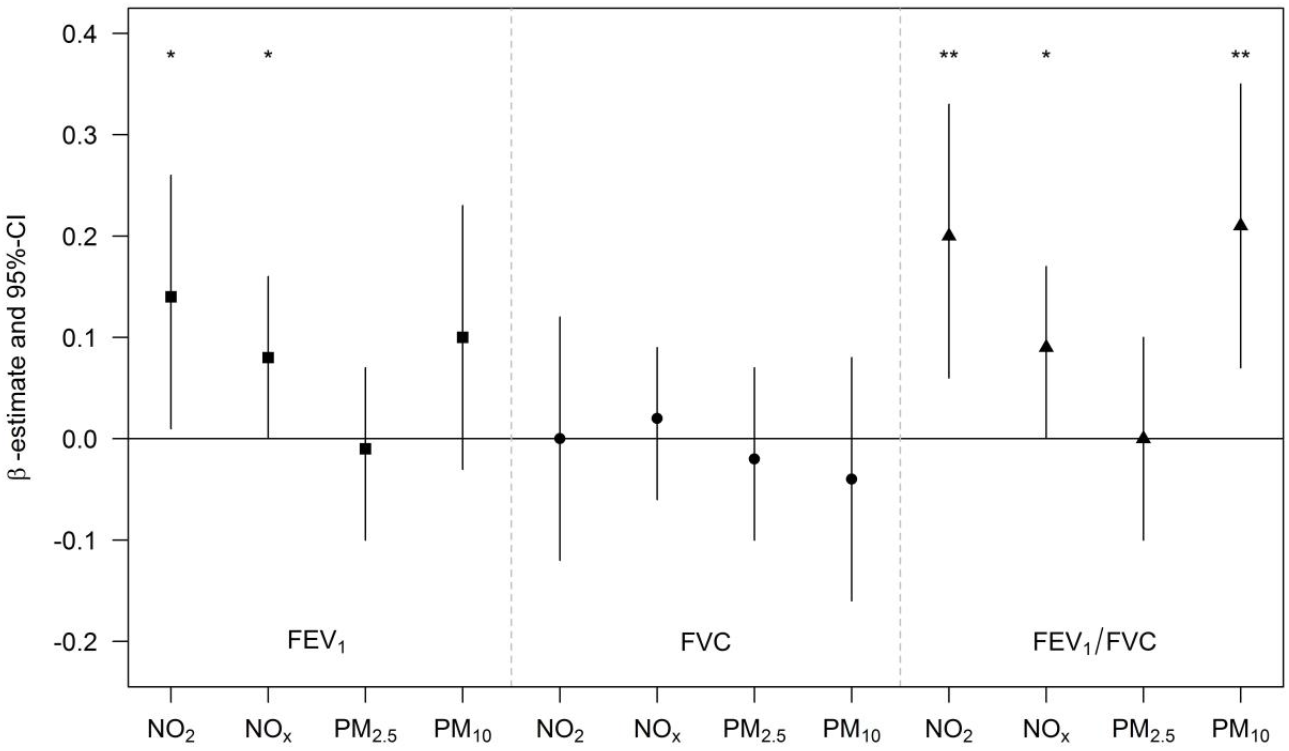
Association between improvement of air quality and change in lung function z-scores over the study period (N=601). β-estimates and 95% confidence intervals (95%-CI) per an improvement of 10 μg/m^3^ in NO_2_, 20 μg/m^3^ in NO_x_, 5 μg/m^3^ in PM_2.5_ and 10 μg/m^3^ in PM_10_. Adjusted for age, BMI at baseline, change of BMI during the study period, level of education, smoking (categorized as current, former or never smoking) and exposure to second hand smoke (SHS). ***: p-value < 0.001; **: 0.001 ≤ p-value < 0.01; *: 0.01 ≤ p-value < 0.05

Associations with NO_2_ and NO_x_ were robust towards adjustment for PM_2.5_, but slightly attenuated after adjustment for PM_10_ (Tables S4). Associations between PM_10_ and FEV_1_/FVC were robust towards adjustment for NO_x_, but not towards adjustment for NO_2_ (Table S5).

Table S6 presents an overview about the lung function related SNPs that were included in the calculation of the GRS (45/49 SNPs available after quality control). None of the single SNPs was associated with a change in lung function after correction for multiple testing.

Combining all SNPs to a GRS, we found a negative interaction between reduced levels of NO_2_ and NO_x_ with the GRS on change in FEV_1_ z-scores (interaction with NO_2_: p=0.029, interaction with NO_x_: p=0.021; Figure 3 and Table S3). These interactions reveal that with an increasing number of lung function related risk alleles, the benefit from improved air quality decreased.

**Figure 3:**
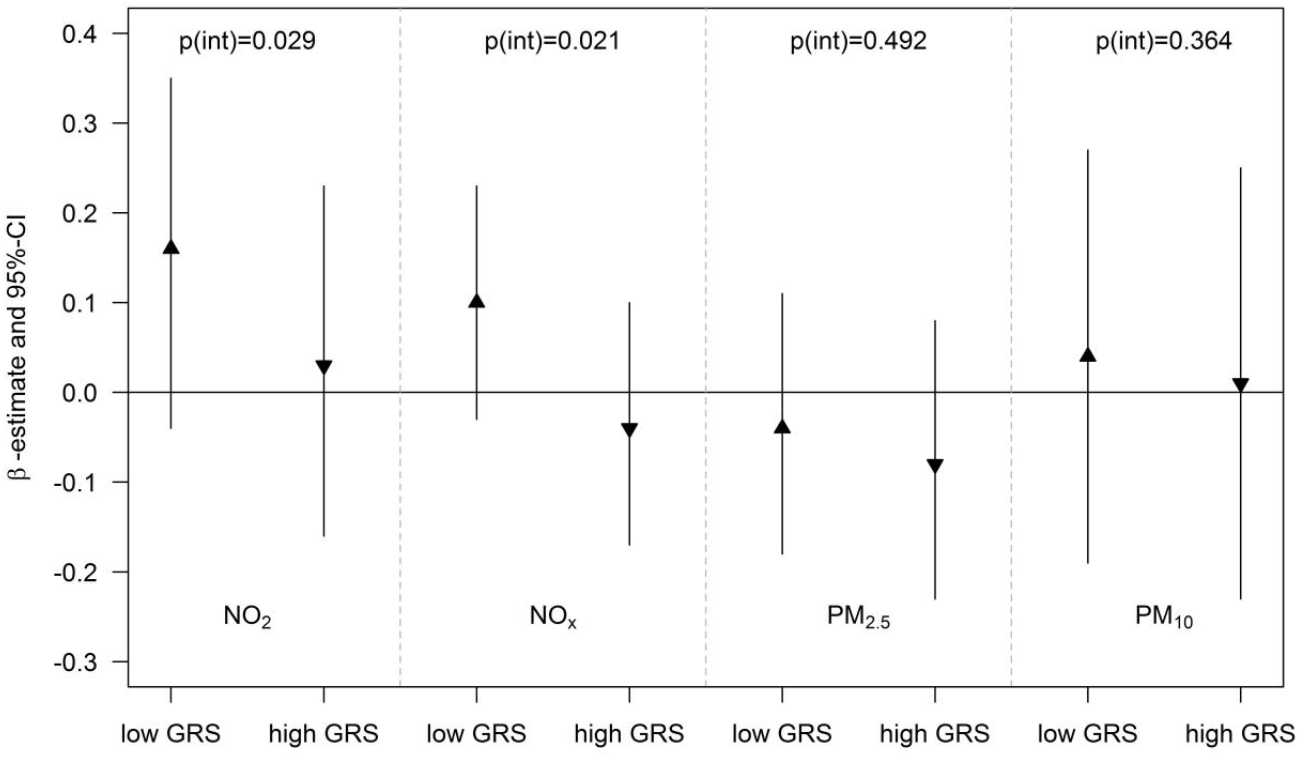
Interaction between GRS of lung function-related SNPs and reduction of air pollution on change in FEV_1_ z-scores over the study period (N=401 with available genotype data). β-estimates and 95% confidence intervals (95%-CI) for the association between improvement of air quality and change in lung function z-scores are given per an improvement of 10 μg/m^3^ in NO_2_, 20 μg/m^3^ in NO_x_, 5 μg/m^3^ in PM_2.5_ and 10 μg/m^3^ in PM_10_ stratified by a low vs. high GRS (cut-point median of GRS). P-values are given for the interaction terms between the continuous GRS and air pollution (p(int)). Adjusted for age, BMI at baseline, change of BMI during the study period, level of education, smoking (categorized as current, former or never smoking) and exposure to second hand smoke (SHS).

Neither the participants’ baseline BMI nor their average BMI during the study period was associated with change in lung function in the elderly (Tables S7 and S8). In the interaction analysis, we found an indication that women with a medium BMI at baseline (BMI of 23.8 to 28.7) or a medium average BMI during the study period (BMI of 25.3 to 28.6) benefit most from improved air quality. However, interactions between BMI and decrease in air pollutants were not significant (Table 3 and Table S9).

**Table 3:** Association between improvement of air quality and change in lung function z-scores stratified by the average BMI within the study period (N=601).

| Change in lung function | Improvement in exposure | 1<sup>st</sup> tertile BMI (<25.3) | 2<sup>nd</sup> tertile BMI (25.3 to 28.6) | 3<sup>rd</sup> tertile BMI (>28.6) | p-value continuous interaction | p-value interaction (1<sup>st</sup> vs. 2<sup>nd</sup>) | p-value interaction (2<sup>nd</sup> vs. 3<sup>rd</sup>) |
| --- | --- | --- | --- | --- | --- | --- | --- |
| FEV<sub>1</sub> | NO<sub>2</sub> (per 10 µg/m<sup>3</sup>) | 0.07 (-0.11; 0.24) | **0.18 (0.02; 0.35)** | 0.13 (-0.04; 0.30) | 0.585 | 0.263 | 0.608 |
|  | NO<sub>x</sub> (per 20 µg/m<sup>3</sup>) | 0.03 (-0.10; 0.16) | **0.12 (0.00; 0.23)** | 0.06 (-0.07; 0.18) | 0.741 | 0.303 | 0.467 |
|  | PM<sub>2.5</sub> (per 5 µg/m<sup>3</sup>) | -0.08 (-0.20; 0.04) | 0.02 (-0.10; 0.14) | -0.02 (-0.15; 0.12) | 0.337 | 0.176 | 0.656 |
|  | PM<sub>10</sub> (per 10 µg/m<sup>3</sup>) | 0.02 (-0.15; 0.19) | 0.18 (0.01; 0.35) | 0.09 (-0.10; 0.28) | 0.369 | 0.128 | 0.433 |
| FVC | NO<sub>2</sub> (per 10 µg/m<sup>3</sup>) | -0.07 (-0.24; 0.10) | 0.08 (-0.08; 0.24) | -0.03 (-0.19; 0.13) | 0.563 | 0.152 | 0.263 |
|  | NO<sub>x</sub> (per 20 µg/m<sup>3</sup>) | -0.01 (-0.14; 0.11) | 0.07 (-0.04; 0.18) | -0.03 (-0.15; 0.09) | 0.745 | 0.305 | 0.216 |
|  | PM<sub>2.5</sub> (per 5 µg/m<sup>3</sup>) | -0.06 (-0.17; 0.06) | 0.03 (-0.08; 0.14) | -0.07 (-0.20; 0.06) | 0.84 | 0.208 | 0.175 |
|  | PM<sub>10</sub> (per 10 µg/m<sup>3</sup>) | -0.09 (-0.25; 0.08) | 0.05 (-0.11; 0.22) | -0.12 (-0.30; 0.07) | 0.882 | 0.156 | 0.110 |
| FEV<sub>1</sub>/FVC | NO<sub>2</sub> (per 10 µg/m<sup>3</sup>) | **0.20 (0.00; 0.40)** | 0.12 (-0.07; 0.31) | **0.26 (0.07; 0.45)** | 0.846 | 0.479 | 0.220 |
|  | NO<sub>x</sub> (per 20 µg/m<sup>3</sup>) | 0.07 (-0.08; 0.21) | 0.05 (-0.08; 0.18) | 0.13 (-0.01; 0.27) | 0.857 | 0.874 | 0.366 |
|  | PM<sub>2.5</sub> (per 5 µg/m<sup>3</sup>) | -0.01 (-0.14; 0.13) | -0.06 (-0.19; 0.07) | 0.09 (-0.06; 0.23) | 0.45 | 0.498 | 0.092 |
|  | PM<sub>10</sub> (per 10 µg/m<sup>3</sup>) | **0.20 (0.01; 0.38)** | 0.13 (-0.06; 0.32) | 0.33 (0.12; 0.54) | 0.378 | 0.56 | 0.104 |

β-estimates and 95% confidence intervals (95%-CI) per an improvement of 10 µg/m<sup>3</sup> in NO<sub>2</sub>, 20 µg/m<sup>3</sup> in NO<sub>x</sub>, 5 µg/m<sup>3</sup> in PM<sub>2.5</sub> and 10 µg/m<sup>3</sup> in PM<sub>10</sub>. P-values are given for the interaction terms between the continuous BMI measurements and air pollution (p-value continuous interaction) as well as for the interaction terms between tertiles of average BMI and air pollution. Adjusted for age, level of education, smoking (categorized as current, former or never smoking) and exposure to second hand smoke (SHS). **Bold:** significant association (p-value < 0.05)

## Discussion

In the present analysis, we used air pollution estimates and spirometric measurements from three time points over a study period of 28 years to show the beneficial effect of improved air quality on lung function in elderly women. In addition, this analysis revealed that the beneficial effects of improved air quality also depended on a person’s genetic make-up. Carriers of more lung function related risk alleles benefited less from improved air quality.

By using data from women up to the age of 83 years, our study extends previous epidemiological studies that showed associations of improved air quality with improvements in childhood lung-function growth [14] as well as with an attenuated decline in lung function among adults [15]. Furthermore, our analysis used data from three measurement time points over a period of 28 years, whereas the maximum duration of previous studies was 11 years with only two measurement time points in adults [15, 18, 19].

In the SALIA study, improved air quality was associated with increased z-scores of FEV_1_ and FEV_1_/FVC, but not with FVC, which is in line with the results of adults from the SAPALDIA cohort in Switzerland [15]. We found the strongest associations for a decrease in NO_2_ and NO_x_ with FEV_1_ as well as with FEV_1_/FVC, whereas a decrease of PM_10_ was only associated with FEV_1_/FVC.

Nitrogen oxides (e.g. NO_2_ and NO_x_) are known to be the best proxy measures of urban-scale variability in chronic exposures to complex urban air pollution mixtures [32]. They are highly correlated with ultrafine particles (UFPs) as well as with black carbon (BC) that both are mainly linked to traffic emissions [32].

Previous studies from the SAPALDIA cohort only included a decrease of PM_10_ as marker of air pollution [12, 15, 18, 19]. Gauderman et al. [14] also included NO_2_, ozone and PM_2.5_ in their study of childhood lung-function growth. However, due to high correlations among reductions of PM_2.5_, PM_10_, and NO_2_, these authors could not assess the independent associations between lung function and each of these pollutants. In contrast, in our study, the correlation between reduction of PM and nitrogen oxides was only moderate, which enabled us to analyze their independent associations with lung function in multi-pollutant models. In these analyses, associations with nitrogen oxides were slightly attenuated after adjustment for PM_10_. Therefore, we could not differentiate the associations with nitrogen oxides from PM.

In the last three decades, air quality has improved substantially in the German Ruhr area. Since the 1970s a series of legislation, e.g., the “lead law”, the Federal Emission Control Act, or administrative regulations such as the Technical Instructions on Air Quality Control and the Large Combustion Installations Ordinance have reduced environmental pollution through technological means. Flue gas desulfurization in power plants, the reduction of sulfur content in all automotive road fuels in the EU, which is now limited to 10 parts per million [33], and less use of coal-fired heating in households have all helped to significantly improve air quality in Germany [34].

However, due to the growth in road traffic in particular vehicles fueled by diesel, NO_2_ levels still exceed the EU limit values in some urban areas. Our observation of improvements in air quality and associated improvements in longitudinal respiratory health provides evidence that changes in traffic-related air pollutant levels can lead to further improved public health.

The beneficial effects of a reduction of air pollution also depend on a person’s genetic make-up. The first GxE interaction analysis on this topic has been conducted in the Swiss SAPALDIA cohort [18], indicating that lung function-associated SNPs slightly modified the association between improvement of air quality and attenuated lung function decline. However, their interaction findings were not robust towards adjustment for multiple testing. Their study differed in two points from our approach. First, Thun et al. [18] only considered ten of the 34 SNPs that had been found to be associated with reduced lung function at the time point of their analysis. In the meantime, the number of significantly associated SNPs has been extended to 49 [16] of which we included 45 in our analysis. Second, Thun et al. [18] conducted a common single SNPs analysis with Bonferroni correction for multiple testing, whereas we conducted a joint SNP analysis, in which we combined all lung function-associated SNPs to a weighted GRS to increase the statistical power to detect interaction effects [29, 35].

In our study of elderly women, there was neither an association between BMI and change in lung function nor an interaction between improvement of air quality and BMI. This is in contrast to Schikowski et al. [19] who found an interaction for improved air quality and change in FVC as well as FEV_1_/FVC in the SAPALDIA cohort, indicating that obese adults might not benefit from improved air quality. The main difference between the two studies is the age range of the participants. In SAPALDIA, the participants had a mean age of 41 years at first follow-up and 52 years at second follow-up, which is still younger than the baseline age in SALIA (55 years). This suggests a different impact of BMI on the beneficial effects of air pollution on lung function during adulthood than at an older age. SAPALDIA further included participants of both sexes, but their interaction findings were not modified by sex [19].

## Strengths & Limitations

The strengths of our study were the long follow-up period of 28 years and the availability of three repeated lung function measures as well as a range of potential confounders prospectively collected over the study period. Furthermore, air pollution exposures were assessed by using state-of-the-art air pollution modelling gained from the ESCAPE campaign in 2009 [23, 24]. In addition, to our knowledge, this is the first longitudinal analysis of air pollution and lung function applying the most recent spirometric reference values (GLI z-scores [27]) that allow to control for age- and height-dependencies in participants up to 93 years of age. Furthermore, we used weighted GRS in our GxE interaction analyses, which is currently considered to be the most powerful approach to detect interactions even in small study populations [29, 35]. However, more studies are needed to replicate our interaction findings.

One limitation of our study is the loss to follow-up, resulting in a reduced study sample at the first and second follow-up investigation. Women lost to follow-up were less well-educated, smoked more heavily, were exposed to higher levels of air pollution and their respiratory health was worse than those who participated in the follow-up [36]. These factors have already been shown to be predictors for cardiovascular mortality in the SALIA cohort [20].

Although exposure estimates were individually assigned to each participant and at each time point of investigation, exposure misclassification is a potential limitation. However, it is unlikely that any unsystematic exposure misclassification had a modifying impact on our results.

## Conclusions

The results of the present analysis suggest a beneficial effect of improved air quality on lung function in elderly women. The elderly are usually considered a high risk group for health effects of air pollution. Thus, these findings support current efforts to further improve air quality. In addition, this study reveals that the beneficial effects of improved air quality depend also on a person’s genetic make-up, which might be helpful for the identification of susceptible subgroups as well as for future treatment strategies.

## Acknowledgments

We thank all study members and staff involved in data collections in each cohort and also the respective funding bodies for SALIA.

Study directorate: R. Dolgner; U. Krämer, U. Ranft, T.Schikowski, A. Vierkötter Scientific Team Baseline: H.W. Schlipköter, M.S. Islam; A. Brockhaus, H. Idel, R. Stiller-Winkler, W. Hadnagy, T. Eikmann,

Scientific Team Follow-up: D. Sugiri, A. Hüls, B. Pesch, A. Hartwig, H. Käfferlein, V. Harth, T.Brüning, T. Weiss

Study Nurses: G. Seitner-Sorge, V. Jäger, G. Petczelies, I. Podolski, T. Hering, M.Goseberg Adminstrative Team: B. Schulten, S. Stolz

*We are most grateful for all the women who participated in the study over decades and the local health Departments for organizing the study.*

Authors’ contribution
U.K, T.S. and B.H. were engaged in the study design of the SALIA study; A.H., U.K., K.I. and T.S. planned the analyses; A.H., D.S. and H.S. prepared and/or analysed the data; A.H. was the major contributor in writing the manuscript; A.H., D.S., M.J.A., B.H., H.S., K.I., U.K. and T.S. revised the manuscript for important intellectual content and approved the final manuscript.

## References

1. Brunekreef B, Beelen R, Hoek G, Schouten L, Bausch-Goldbohm S, Fischer P, Armstrong B, Hughes E, Jerrett M, van den Brandt P. Effects of long-term exposure to traffic-related air pollution on respiratory and cardiovascular mortality in the Netherlands: the NLCS-AIR study. Res. Rep. Health. Eff. Inst. 2009;: 5–71; discussion 73-89.

2. Beelen R, Raaschou-Nielsen O, Stafoggia M, Andersen ZJ, Weinmayr G, Hoffmann B, Wolf K, Samoli E, Fischer P, Nieuwenhuijsen M, Vineis P, Xun WW, Katsouyanni K, Dimakopoulou K, Oudin A, Forsberg B, Modig L, Havulinna AS, Lanki T, Turunen A, Oftedal B, Nystad W, Nafstad P, De Faire U, Pedersen NL, Östenson C-G, Fratiglioni L, Penell J, Korek M, Pershagen G, et al. Effects of long-term exposure to air pollution on natural-cause mortality: an analysis of 22 European cohorts within the multicentre ESCAPE project. Lancet 2014; 383: 785–795.

3. Schikowski T, Sugiri D, Ranft U, Gehring U, Heinrich J, Wichmann H-E, Krämer U. Long-term air pollution exposure and living close to busy roads are associated with COPD in women. Respir. Res. 2005; 6: 152.

4. Strak M, Boogaard H, Meliefste K, Oldenwening M, Zuurbier M, Brunekreef B, Hoek G. Respiratory health effects of ultrafine and fine particle exposure in cyclists. Occup. Environ. Med. 2010; 67: 118–124.

5. Cesaroni G, Badaloni C, Porta D, Forastiere F, Perucci CA. Comparison between various indices of exposure to traffic-related air pollution and their impact on respiratory health in adults. Occup. Environ. Med. 2008; 65: 683–690.

6. Ackermann-Liebrich U, Leuenberger P, Schwartz J, Schindler C, Monn C, Bolognini G, Bongard JP, Brändli O, Domenighetti G, Elsasser S, Grize L, Karrer W, Keller R, Keller-Wossidlo H, Künzli N, Martin BW, Medici TC, Perruchoud AP, Schöni MH, Tschopp JM, Villiger B, Wüthrich B, Zellweger JP, Zemp E. Lung function and long term exposure to air pollutants in Switzerland. Study on Air Pollution and Lung Diseases in Adults (SAPALDIA) Team. Am. J. Respir. Crit. Care Med. 1997; 155: 122–129.

7. Brunekreef B, Holgate ST. Air pollution and health. Lancet 2002; 360: 1233–1242.

8. Suglia SF, Gryparis A, Schwartz J, Wright RJ. Association between traffic-related black carbon exposure and lung function among urban women. Environ. Health Perspect. 2008; 116: 1333–1337.

9. Götschi T, Heinrich J, Sunyer J, Künzli N. Long-term effects of ambient air pollution on lung function: a review. Epidemiology 2008; 19: 690–701.

10. Kan H, Heiss G, Rose KM, Whitsel E, Lurmann F, London SJ. Traffic exposure and lung function in adults: the Atherosclerosis Risk in Communities study. Thorax 2007; 62: 873–879.

11. Adam M, Schikowski T, Carsin AE, Cai Y, Jacquemin B, Sanchez M, Vierkötter A, Marcon A, Keidel D, Sugiri D, Al Kanani Z, Nadif R, Siroux V, Hardy R, Kuh D, Rochat T, Bridevaux P-O, Eeftens M, Tsai M-Y, Villani S, Phuleria HC, Birk M, Cyrys J, Cirach M, de Nazelle A, Nieuwenhuijsen MJ, Forsberg B, de Hoogh K, Declerq C, Bono R, et al. Adult lung function and long-term air pollution exposure. ESCAPE: a multicentre cohort study and meta-analysis. Eur. Respir. J. 2015; 45: 38–50.

12. Schindler C, Keidel D, Gerbase MW, Zemp E, Bettschart R, Brändli O, Brutsche MH, Burdet L, Karrer W, Knöpfli B, Pons M, Rapp R, Bayer-Oglesby L, Künzli N, Schwartz J, Liu L-JS, Ackermann-Liebrich U, Rochat T. Improvements in PM10 exposure and reduced rates of respiratory symptoms in a cohort of Swiss adults (SAPALDIA). Am. J. Respir. Crit. Care Med. [Internet] 2009; 179: 579–587 Available from: http://www.ncbi.nlm.nih.gov/pubmed/19151198.

13. Schikowski T, Ranft U, Sugiri D, Vierkötter A, Brüning T, Harth V, Krämer U. Decline in air pollution and change in prevalence in respiratory symptoms and chronic obstructive pulmonary disease in elderly women. Respir. Res. 2010; 11: 113.

14. Gauderman WJ, Urman R, Avol E, Berhane K, McConnell R, Rappaport E, Chang R, Lurmann F, Gilliland F. Association of improved air quality with lung development in children. N. Engl. J. Med. 2015; 372: 905–913.

15. Downs SH, Schindler C, Liu L-JS, Keidel D, Bayer-Oglesby L, Brutsche MH, Gerbase MW, Keller R, Künzli N, Leuenberger P, Probst-Hensch NM, Tschopp J-M, Zellweger J-P, Rochat T, Schwartz J, Ackermann-Liebrich U. Reduced exposure to PM10 and attenuated age-related decline in lung function. N. Engl. J. Med. 2007; 357: 2338–2347.

16. Soler Artigas M, Wain L V, Miller S, Kheirallah AK, Huffman JE, Ntalla I, Shrine N, Obeidat M, Trochet H, McArdle WL, Alves AC, Hui J, Zhao JH, Joshi PK, Teumer A, Albrecht E, Imboden M, Rawal R, Lopez LM, Marten J, Enroth S, Surakka I, Polasek O, Lyytikäinen L-P, Granell R, Hysi PG, Flexeder C, Mahajan A, Beilby J, Bossé Y, et al. Sixteen new lung function signals identified through 1000 Genomes Project reference panel imputation. Nat. Commun. 2015; 6: 8658.

17. Leone N, Courbon D, Thomas F, Bean K, Jégo B, Leynaert B, Guize L, Zureik M. Lung function impairment and metabolic syndrome. The critical role of abdominal obesity. Am. J. Respir. Crit. Care Med. 2009; 179: 509–516.

18. Thun GA, Imboden M, Künzli N, Rochat T, Keidel D, Haun M, Schindler C, Kronenberg F, Probst-Hensch NM. Follow-up on genome-wide main effects: Do polymorphisms modify the air pollution effect on lung function decline in adults? Environ. Int. 2014; 64: 110–115.

19. Schikowski T, Schaffner E, Meier F, Phuleria HC, Vierkötter A, Schindler C, Kriemler S, Zemp E, Krämer U, Bridevaux P-O, Rochat T, Schwartz J, Künzli N, Probst-Hensch N. Improved air quality and attenuated lung function decline: modification by obesity in the SAPALDIA cohort. Environ. Health Perspect. 2013; 121: 1034–1039.

20. Schikowski T, Sugiri D, Ranft U, Gehring U, Heinrich J, Wichmann H-E, Krämer U. Does respiratory health contribute to the effects of long-term air pollution exposure on cardiovascular mortality? Respir. Res. 2007; 8: 20.

21. Schikowski T, Vossoughi M, Vierkotter A, Schulte T, Teichert T, Sugiri D, Fehsel K, Tzivian L, Bae I-S, Ranft U, Hoffmann B, Probst-Hensch N, Herder C, Kramer U, Luckhaus C. Association of air pollution with cognitive functions and its modification by APOE gene variants in elderly women. Environ. Res. Elsevier; 2015; 142: 10–16.

22. Hüls A, Krämer U, Stolz S, Hennig F, Hoffmann B, Ickstadt K, Vierkötter A, Schikowski T. Applicability of the Global Lung Initiative 2012 Reference Values for Spirometry for Longitudinal Data of Elderly Women. PLoS One 2016; 11.

23. Eeftens M, Beelen R, de Hoogh K, Bellander T, Cesaroni G, Cirach M, Declercq C, Dėdelė A, Dons E, de Nazelle A, Dimakopoulou K, Eriksen K, Falq G, Fischer P, Galassi C, Gražulevičienė R, Heinrich J, Hoffmann B, Jerrett M, Keidel D, Korek M, Lanki T, Lindley S, Madsen C, Mölter A, Nádor G, Nieuwenhuijsen M, Nonnemacher M, Pedeli X, Raaschou-Nielsen O, et al. Development of Land Use Regression Models for PM 2.5, PM 2.5 Absorbance, PM 10 and PM coarse in 20 European Study Areas; Results of the ESCAPE Project. Environ. Sci. Technol. American Chemical Society; 2012; 46: 11195–11205.

24. Beelen R, Hoek G, Vienneau D, Eeftens M, Dimakopoulou K, Pedeli X, Tsai M-Y, Künzli N, Schikowski T, Marcon A, Eriksen KT, Raaschou-Nielsen O, Stephanou E, Patelarou E, Lanki T, Yli-Tuomi T, Declercq C, Falq G, Stempfelet M, Birk M, Cyrys J, von Klot S, Nádor G, Varró MJ, Dėdelė A, Gražulevičienė R, Mölter A, Lindley S, Madsen C, Cesaroni G, et al. Development of NO2 and NOx land use regression models for estimating air pollution exposure in 36 study areas in Europe – The ESCAPE project. Atmos. Environ. 2013; 72: 10–23.

25. Das S, Forer L, Schönherr S, Sidore C, Locke AE, Kwong A, Vrieze SI, Chew EY, Levy S, McGue M, Schlessinger D, Stambolian D, Loh PR, Iacono WG, Swaroop A, Scott LJ, Cucca F, Kronenberg F, Boehnke M, Abecasis GR, Fuchsberger C. Next-generation genotype imputation service and methods. Nat. Genet. 2016; 48: 1284–1287.

26. Miller MR, Hankinson J, Brusasco V, Burgos F, Casaburi R, Coates A, Crapo R, Enright P, van der Grinten CPM, Gustafsson P, Jensen R, Johnson DC, MacIntyre N, McKay R, Navajas D, Pedersen OF, Pellegrino R, Viegi G, Wanger J. Standardisation of spirometry. Eur. Respir. J. 2005; 26: 319–338.

27. Quanjer PH, Stanojevic S, Cole TJ, Baur X, Hall GL, Culver BH, Enright PL, Hankinson JL, Ip MSM, Zheng J, Stocks J. Multi-ethnic reference values for spirometry for the 3-95-yr age range: the global lung function 2012 equations. Eur. Respir. J. 2012; 40: 1324–1343.

28. Twisk JWR. Applied Longitudinal Data Analysis for Epidemiology: A Practical Guide. Cambridge University Press; 2007.

29. Hüls A, Ickstadt K, Schikowski T, Krämer U. Detection of gene-environment interactions in the presence of linkage disequilibrium and noise by using genetic risk scores with internal weights from elastic net regression. BMC Genet. 2017; 18: 55.

30. Hüls A, Krämer U, Carlsten C, Schikowski T, Ickstadt K, Schwender H. Comparison of weighting approaches for genetic risk scores in gene-environment interaction studies. BMC Genet. 2017; 18: 115.

31. R Development Core Team. R: A language and environment for statistical computing [Internet]. Vienna, Austria: R Foundation for Statistical Computing; 2018. Available from: http://www.r-project.org/.

32. Levy I, Mihele C, Lu G, Narayan J, Brook JR. Evaluating multipollutant exposure and urban air quality: Pollutant interrelationships, neighborhood variability, and nitrogen dioxide as a proxy pollutant. Environ. Health Perspect. 2014; 122: 65–72.

33. Mellios G, Kouridis C. EU Fuel Quality Monitoring – 2014 Summary Report. European Environment Agency (EEA) Technical report. 2015.

34. Flasbarth J. Federal Environment Agency: The sky over the Ruhr is blue again! [Internet]. 25/2011 [cited 2018 Apr 3]. Available from: https://www.umweltbundesamt.de/en/press/pressinformation/federal-environment-agency-sky-over-ruhr-is-blue.

35. Aschard H. A perspective on interaction effects in genetic association studies. Genet. Epidemiol. 2016; 40: 678–688.

36. Hüls A, Vierkötter A, Sugiri D, Abramson M, Ranft U, Krämer U, Schikowski T. The role of air pollution and lung function on cognitive impairment. Eur. Respir. J. 2018; 51: 1701963.

