## Supplementary Material for "Benefits of improved air quality on aging lungs – Impacts of genetics and obesity"

### *Methods*

#### *Assessment of air pollution*

Outdoor air pollution concentrations were assessed according to the ESCAPE (European Study of Cohorts for Air Pollution Effects) protocol [1, 2]. Each participant was assigned an annual average concentration of $NO_2$, $PM_{2.5}$ and $PM_{10}$ at home. Air pollution was monitored over one year (2008-2009) in three two-week periods in the Ruhr area and the adjacent Muensterland at overall 40 monitoring sites. Three two-week measurements were performed at 40 (20 for PM) monitoring sites, one measurement in cold, warm and intermediate season, and adjusted for temporal variation by a reference station, which measured during the whole period. The measured concentrations at the monitoring sites were associated with characteristics of land-use at this location to obtain a regression equation (land-use regression (LUR) model). Data on nearby traffic, ports or industry and population/household density derived from Geographic Information Systems (GIS) were included in the equations. The models were applied to the home addresses to get individual exposure concentrations.

To characterize the levels of air pollution exposure at each time point of investigation, we used extrapolation procedures based on air pollution measurements from the dedicated ESCAPE monitoring campaign in 2008/2009 in the area of SALIA residential addresses. Ratios were calculated between air pollution concentrations at government background monitoring stations for $PM_{10}$ (and additionally for $PM_{2.5}$ in 2012/2013) and $NO_2$ during a two-year period around each time point of investigation and the period of ESCAPE monitoring campaign. The values from the LUR models were corrected for spatial trends by multiplying them with these ratios. The implicit assumption of proportional spatial contrasts over time was validated with data from six routine monitoring stations situated in the study area and covering the investigation period.

#### *Assessment of genotypes*

A genome-wide genotyping was performed in 462 women who participated in the follow-up investigation 2007-2010, using the Affymetrix Axiom™ Precision Medicine Research Array. SNPs were imputed on the 1000 Genomes reference panel (Phase III) using Minimac3 [3]. We considered only SNPs with a minor allele frequency >1% and a moderate imputation quality score ($R^2$>0.3).

*Assessment of potential confounders*

Information was collected on age, measured BMI, socioeconomic status, current and past smoking habits, passive smoking exposure at home or work, and other risk factors. We classified socioeconomic status at baseline into three categories using the highest school level achieved by either the woman or her husband as low (less than 10 years), medium (=10 years) or high (more than 10 years).

**Table S1: Complete cases.** Description of study population at baseline, first and second follow-up (including lung function). All women with lung function measurements at three time points of measurement were included in the analysis (N=319).

| | **Baseline (1985-1994)** | **1st Follow-up (2007-2010)** | **2nd Follow-up (2012-2013)** |
|---|---|---|---|
| N | 319 | 319 | 319 |
| Age years, mean ± sd | 54.3 +- 0.8 | 73.2 +- 3.3 | 77.3 +- 3.5 |
| BMI kg/m$^2$, mean ± sd | 26.4 +- 3.7 | 27.1 +- 4.3 | 28.4 +- 4.6 |
| Change from baseline BMI*, mean ± sd | | 0.8 +- 2.7 | 2.0 +- 3.0 |
| >10 years of education, n (%) | 114 (35.7%) | 114 (35.7%) | 114 (35.7%) |
| 10 years of education, n (%) | 140 (43.9%) | 140 (43.9%) | 140 (43.9%) |
| <10 years of education, n (%) | 65 (20.4%) | 65 (20.4%) | 65 (20.4%) |
| Smoker, n (%) | 28 (8.8%) | 12 (3.8%) | 12 (3.8%) |
| Ex-smoker, n (%) | 30 (9.4%) | 48 (15.0%) | 47 (14.7%) |
| Never smoker, n (%) | 260 (81.5%) | 259 (81.2%) | 260 (81.5%) |
| Passive smoking, n (%) | 147 (46.1%) | 187 (58.6%) | 197 (61.8%) |
| GLI z-score $FEV_1$, mean ± sd | -0.2 +- 1.0 | 0.3 +- 1.0 | 0.1 +- 1.1 |
| Change from baseline GLI z-score $FEV_1$[†], mean ± sd | | 0.4 +- 0.8 | 0.2 +- 0.8 |
| GLI z-score FVC, mean ± sd | 0.0 +- 0.9 | 0.3 +- 0.9 | 0.5 +- 1.0 |
| Change from baseline GLI z-score FVC[†], mean ± sd | | 0.3 +- 0.7 | 0.4 +- 0.8 |
| GLI z-score $FEV_1$/FVC, mean ± sd | -0.4 +- 0.8 | -0.2 +- 0.9 | -0.7 +- 0.8 |
| Change from baseline GLI z-score $FEV_1$/FVC[†], mean ± sd | | 0.2 +- 0.9 | -0.3 +- 0.8 |

*Change of BMI between baseline and 1st follow-up (BMI at 1st follow-up - BMI at baseline) and between baseline and 2nd follow-up (BMI at 2nd follow-up – BMI at baseline).
[†]Change of lung function between baseline and 1st follow-up (lung function at 1st follow-up - lung function at baseline) and between baseline and 2nd follow-up (lung function at 2nd follow-up – lung function at baseline).

**Table S2:** Pearson correlations between reduction of different air pollutants (in µg/m$^3$) between baseline (B) and first follow-up (F1) and between baseline (B) and second follow-up (F2) (N=601).

| | **$NO_2$ from B to F1** | **$NO_x$ from B to F1** | **$PM_{2.5}$ from B to F1** | **$PM_{10}$ from B to F1** | **$NO_2$ from B to F2** | **$NO_x$ from B to F2** | **$PM_{2.5}$ from B to F2** | **$PM_{10}$ from B to F2** |
|---|---|---|---|---|---|---|---|---|
| **$NO_2$ from B to F1** | 1 | 0.88 | 0.69 | 0.72 | 0.97 | 0.86 | 0.73 | 0.75 |
| **$NO_x$ from B to F1** | | 1 | 0.66 | 0.63 | 0.88 | 0.98 | 0.65 | 0.63 |
| **$PM_{2.5}$ from B to F1** | | | 1 | 0.98 | 0.7 | 0.65 | 0.98 | 0.98 |
| **$PM_{10}$ from B to F1** | | | | 1 | 0.74 | 0.62 | 0.98 | 0.99 |
| **$NO_2$ from B to F2** | | | | | 1 | 0.9 | 0.73 | 0.75 |
| **$NO_x$ from B to F2** | | | | | | 1 | 0.66 | 0.64 |
| **$PM_{2.5}$ from B to F2** | | | | | | | 1 | 0.99 |
| **$PM_{10}$ from B to F2** | | | | | | | | 1 |

**Table S3:** Association between improvement of air quality and change in lung function z-scores in the whole study population and stratified by carrying few lung function related risk alleles (low GRS) vs. carrying many lung function related risk alleles (high GRS; cut-point median of GRS).

| **Change in lung function** | **Improvement in exposure** | **All*** | **Low GRS†** | **High GRS†** | **p-value continous interaction†** |
|---|---|---|---|---|---|
| $FEV_1$ | $NO_2$ (per 10 µg/m$^3$) | **0.14 (0.01; 0.26)** | 0.16 (-0.04; 0.35) | 0.03 (-0.16; 0.23) | **0.029** |
| | $NO_x$ (per 20 µg/m$^3$) | **0.08 (0.00; 0.16)** | 0.10 (-0.03; 0.23) | -0.04 (-0.17; 0.10) | **0.021** |
| | $PM_{2.5}$ (per 5 µg/m$^3$) | -0.01 (-0.10; 0.07) | -0.04 (-0.18; 0.11) | -0.08 (-0.23; 0.08) | 0.492 |
| | $PM_{10}$ (per 10 µg/m$^3$) | 0.10 (-0.03; 0.23) | 0.04 (-0.19; 0.27) | 0.01 (-0.23; 0.25) | 0.364 |
| FVC | $NO_2$ (per 10 µg/m$^3$) | 0.00 (-0.12; 0.12) | 0.07 (-0.12; 0.26) | -0.01 (-0.20; 0.18) | 0.227 |
| | $NO_x$ (per 20 µg/m$^3$) | 0.02 (-0.06; 0.09) | 0.09 (-0.03; 0.21) | -0.03 (-0.16; 0.10) | 0.217 |
| | $PM_{2.5}$ (per 5 µg/m$^3$) | -0.02 (-0.10; 0.07) | 0.03 (-0.11; 0.17) | 0.06 (-0.09; 0.21) | 0.855 |
| | $PM_{10}$ (per 10 µg/m$^3$) | -0.04 (-0.16; 0.08) | 0.01 (-0.21; 0.23) | 0.10 (-0.12; 0.33) | 0.997 |
| $FEV_1$/FVC | $NO_2$ (per 10 µg/m$^3$) | **0.20 (0.06; 0.33)** | 0.08 (-0.13; 0.30) | 0.08 (-0.14; 0.29) | 0.273 |
| | $NO_x$ (per 20 µg/m$^3$) | **0.09 (0.00; 0.17)** | -0.03 (-0.17; 0.11) | -0.01 (-0.16; 0.14) | 0.210 |
| | $PM_{2.5}$ (per 5 µg/m$^3$) | 0.00 (-0.10; 0.10) | -0.16 (-0.32; 0.00) | -0.21 (-0.38; -0.04) | 0.350 |
| | $PM_{10}$ (per 10 µg/m$^3$) | **0.21 (0.07; 0.35)** | 0.02 (-0.24; 0.27) | -0.10 (-0.37; 0.16) | 0.339 |

β-estimates and 95% confidence intervals (95%-CI) per an improvement of 10 µg/m$^3$ in $NO_2$, 20 µg/m$^3$ in $NO_x$, 5 µg/m$^3$ in $PM_{2.5}$ and 10 µg/m$^3$ in $PM_{10}$. P-values are given for the interaction terms between the continuous GRS and air pollution. Adjusted for age, BMI at baseline, change of BMI during the study period, level of education, smoking (categorized as current, former or never smoking) and exposure to second hand smoke (SHS). **Bold**: significant association or interaction (p-value < 0.05)

*All women with lung function measurements at ≥2 time points of measurement (N=601); †subset with available genotype data (N=401)

**Table S4: Multi-pollutant models.** Association between improvement of nitrogen oxides ($NO_2$ and $NO_x$) and change in lung function z-scores in one- and two-pollutant models (adjusted for $PM_{2.5}$ and $PM_{10}$).

| Change in lung function | Improvement in exposure | One-pollutant model | Adjusted for $PM_{2.5}$ | Adjusted for $PM_{10}$ |
|---|---|---|---|---|
| $FEV_1$ | $NO_2$ (per 10 µg/m$^3$) | **0.14 (0.01; 0.26)** | **0.19 (0.05; 0.33)** | 0.12 (-0.02; 0.26) |
| | $NO_x$ (per 20 µg/m$^3$) | **0.08 (0.00; 0.16)** | **0.13 (0.04; 0.23)** | 0.07 (-0.02; 0.16) |
| FVC | $NO_2$ (per 10 µg/m$^3$) | 0.00 (-0.12; 0.12) | 0.02 (-0.11; 0.15) | 0.03 (-0.11; 0.16) |
| | $NO_x$ (per 20 µg/m$^3$) | 0.02 (-0.06; 0.09) | 0.04 (-0.05; 0.13) | 0.05 (-0.04; 0.13) |
| $FEV_1$/FVC | $NO_2$ (per 10 µg/m$^3$) | **0.20 (0.06; 0.33)** | **0.26 (0.10; 0.41)** | 0.13 (-0.03; 0.29) |
| | $NO_x$ (per 20 µg/m$^3$) | **0.09 (0.00; 0.17)** | **0.13 (0.02; 0.24)** | 0.03 (-0.08; 0.13) |

β-estimates and 95% confidence intervals (95%-CI) per an improvement of 10 µg/m$^3$ in $NO_2$, 20 µg/m$^3$ in $NO_x$, 5 µg/m$^3$ in $PM_{2.5}$ and 10 µg/m$^3$ in $PM_{10}$. Adjusted for age, BMI at baseline, change of BMI during the study period, level of education, smoking (categorized as current, former or never smoking) and exposure to second hand smoke (SHS). **Bold**: significant association ($p$-value < 0.05)

All women with lung function measurements at ≥2 time points of measurement (N=601)

**Table S5: Multi-pollutant models.** Association between improvement of PM ($PM_{2.5}$ and $PM_{10}NO_2$ and $NO_x$) and change in lung function z-scores in one- and two-pollutant models (adjusted for $NO_2$ and $NO_x$).

| Change in lung function | Improvement in exposure | One-pollutant model | Adjusted for $NO_2$ | Adjusted for $NO_x$ |
|---|---|---|---|---|
| $FEV_1$ | $PM_{2.5}$ (per 5 µg/m$^3$) | -0.01 (-0.10; 0.07) | -0.08 (-0.18; 0.02) | -0.10 (-0.21; 0.01) |
| | $PM_{10}$ (per 10 µg/m$^3$) | 0.10 (-0.03; 0.23) | 0.04 (-0.11; 0.19) | 0.04 (-0.11; 0.19) |
| FVC | $PM_{2.5}$ (per 5 µg/m$^3$) | -0.02 (-0.10; 0.07) | -0.03 (-0.12; 0.07) | -0.05 (-0.15; 0.06) |
| | $PM_{10}$ (per 10 µg/m$^3$) | -0.04 (-0.16; 0.08) | -0.06 (-0.20; 0.08) | -0.08 (-0.23; 0.06) |
| $FEV_1$/FVC | $PM_{2.5}$ (per 5 µg/m$^3$) | 0.00 (-0.10; 0.10) | -0.09 (-0.20; 0.02) | -0.08 (-0.20; 0.04) |
| | $PM_{10}$ (per 10 µg/m$^3$) | **0.21 (0.07; 0.35)** | 0.14 (-0.02; 0.31) | **0.19 (0.02; 0.36)** |

β-estimates and 95% confidence intervals (95%-CI) per an improvement of 10 µg/m$^3$ in $NO_2$, 20 µg/m$^3$ in $NO_x$, 5 µg/m$^3$ in $PM_{2.5}$ and 10 µg/m$^3$ in $PM_{10}$. Adjusted for age, BMI at baseline, change of BMI during the study period, level of education, smoking (categorized as current, former or never smoking) and exposure to second hand smoke (SHS). **Bold**: significant association (p-value < 0.05)

All women with lung function measurements at ≥2 time points of measurement (N=601)

**Table S6:** Overview about SNPs that were shown to be associated with impaired lung function in genome-wide association studies (GWAS) [4] and available in the SALIA cohort. Associations between effect alleles (additive model) and change in GLI z-scores for $FEV_1$, FVC and $FEV_1$/FVC over the study period.

| | | | | | | | **$FEV_1$** | | **FVC** | | **$FEV_1$/FVC** | |
|---|---|---|---|---|---|---|---|---|---|---|---|---|
| **Chr** | **Pos** | **rsID** | **Genotyped / Imputed** | **$R^2$** | **EA/OA** | **EA Freq** | **Estimate** | **p-value** | **Estimate** | **p-value** | **Estimate** | **p-value** |
| 1 | 17306675 | rs2284746 | Genotyped | 1.00 | G/C | 0.56 | 0.02 | 0.705 | 0.03 | 0.524 | -0.06 | 0.359 |
| 1 | 150586971 | rs6681426 | Imputed | 0.79 | A/G | 0.65 | -0.01 | 0.876 | 0.02 | 0.776 | -0.04 | 0.546 |
| 1 | 218860068 | rs993925 | Genotyped | 1.00 | T/C | 0.33 | -0.05 | 0.408 | -0.05 | 0.314 | 0.01 | 0.851 |
| 2 | 18292452 | rs61067109 | Imputed | 0.99 | A/G | 0.24 | 0.05 | 0.452 | -0.01 | 0.910 | 0.09 | 0.205 |
| 2 | 56120853 | rs1430193 | Genotyped | 1.00 | T/A | 0.37 | **-0.12** | **0.042** | -0.03 | 0.554 | **-0.13** | **0.047** |
| 2 | 218683154 | rs2571445 | Genotyped | 1.00 | G/A | 0.59 | 0.06 | 0.318 | 0.05 | 0.388 | 0.00 | 0.992 |
| 2 | 239877148 | rs12477314 | Genotyped | 1.00 | T/C | 0.20 | -0.05 | 0.520 | -0.04 | 0.516 | 0.02 | 0.852 |
| 3 | 25520582 | rs1529672 | Imputed | 0.89 | A/C | 0.16 | 0.12 | 0.122 | 0.05 | 0.492 | 0.08 | 0.341 |
| 3 | 158282459 | rs6441207 | Imputed | 0.93 | T/C | 0.42 | 0.10 | 0.068 | 0.08 | 0.129 | 0.01 | 0.827 |
| 3 | 169300219 | rs1344555 | Genotyped | 1.00 | T/C | 0.21 | 0.05 | 0.441 | 0.12 | 0.062 | -0.10 | 0.185 |
| 4 | 89870964 | rs2045517 | Imputed | 1.00 | T/C | 0.39 | -0.01 | 0.880 | 0.01 | 0.863 | -0.03 | 0.685 |
| 4 | 106688904 | rs10516526 | Genotyped | 1.00 | G/A | 0.06 | -0.06 | 0.615 | -0.04 | 0.711 | 0.00 | 0.985 |
| 4 | 106841962 | rs6856422 | Imputed | 0.86 | T/G | 0.40 | 0.04 | 0.529 | 0.04 | 0.402 | 0.03 | 0.649 |
| 4 | 145479139 | rs11100860 | Imputed | 1.00 | G/A | 0.46 | -0.02 | 0.785 | 0.03 | 0.557 | -0.10 | 0.104 |
| 5 | 95036700 | rs153916 | Genotyped | 1.00 | T/C | 0.55 | -0.02 | 0.766 | -0.02 | 0.701 | 0.02 | 0.803 |
| 5 | 147847788 | rs1985524 | Imputed | 0.94 | C/G | 0.44 | 0.04 | 0.478 | 0.00 | 0.957 | 0.06 | 0.327 |
| 5 | 156936766 | rs11134779 | Imputed | 0.99 | G/A | 0.32 | 0.04 | 0.551 | 0.09 | 0.129 | -0.07 | 0.299 |

| | | | | | | | | | | | | |
|---|---|---|---|---|---|---|---|---|---|---|---|---|
| 6 | 7801112 | rs6923462 | Genotyped | 1.00 | C/T | 0.15 | 0.02 | 0.812 | 0.00 | 0.966 | 0.02 | 0.809 |
| 6 | 28322296 | rs6903823 | Genotyped | 1.00 | G/A | 0.22 | 0.00 | 0.971 | -0.07 | 0.391 | 0.09 | 0.345 |
| 6 | 31568469 | rs2857595 | Genotyped | 1.00 | A/G | 0.21 | -0.09 | 0.296 | -0.09 | 0.294 | -0.05 | 0.614 |
| 6 | 32151443 | rs2070600 | Genotyped | 1.00 | T/C | 0.05 | 0.12 | 0.385 | 0.16 | 0.210 | -0.07 | 0.621 |
| 6 | 109268050 | rs2798641 | Genotyped | 1.00 | T/C | 0.21 | **-0.15** | **0.032** | -0.03 | 0.645 | **-0.17** | **0.026** |
| 6 | 142838173 | rs148274477 | Imputed | 0.56 | T/C | 0.02 | -0.05 | 0.802 | 0.00 | 0.994 | -0.12 | 0.602 |
| 6 | 142853144 | rs262129 | Imputed | 0.75 | G/A | 0.31 | -0.05 | 0.402 | -0.09 | 0.093 | 0.07 | 0.310 |
| 9 | 98204792 | rs16909859 | Imputed | 0.92 | A/G | 0.07 | 0.02 | 0.846 | -0.15 | 0.151 | **0.26** | **0.037** |
| 9 | 98231008 | rs16909898 | Genotyped | 1.00 | G/A | 0.09 | -0.01 | 0.938 | -0.09 | 0.339 | 0.12 | 0.261 |
| 9 | 139094805 | rs2274116 | Imputed | 0.96 | T/C | 0.33 | 0.08 | 0.191 | 0.03 | 0.607 | 0.06 | 0.386 |
| 10 | 12277992 | rs7068966 | Genotyped | 1.00 | T/C | 0.54 | -0.02 | 0.753 | 0.04 | 0.414 | -0.07 | 0.271 |
| 10 | 78315224 | rs11001819 | Genotyped | 1.00 | A/G | 0.48 | 0.03 | 0.643 | -0.01 | 0.782 | 0.05 | 0.376 |
| 11 | 43648368 | rs4237643 | Imputed | 0.98 | G/T | 0.70 | 0.01 | 0.912 | 0.01 | 0.823 | 0.01 | 0.879 |
| 11 | 45250732 | rs2863171 | Genotyped | 1.00 | C/A | 0.14 | 0.01 | 0.916 | -0.08 | 0.334 | 0.14 | 0.137 |
| 12 | 57527283 | rs11172113 | Genotyped | 0.99 | C/T | 0.44 | 0.06 | 0.294 | 0.03 | 0.547 | 0.07 | 0.284 |
| 12 | 96271428 | rs1036429 | Genotyped | 0.99 | C/T | 0.78 | 0.10 | 0.166 | 0.12 | 0.070 | -0.02 | 0.835 |
| 12 | 115201436 | rs10850377 | Imputed | 0.91 | A/G | 0.34 | -0.03 | 0.660 | -0.04 | 0.504 | 0.04 | 0.582 |
| 14 | 92485881 | rs7155279 | Genotyped | 1.00 | T/G | 0.38 | 0.05 | 0.384 | 0.02 | 0.654 | 0.06 | 0.365 |
| 14 | 93118229 | rs117068593 | Imputed | 0.93 | T/C | 0.15 | 0.12 | 0.130 | 0.04 | 0.560 | 0.11 | 0.193 |
| 15 | 71680080 | rs8033889 | Imputed | 0.97 | T/G | 0.19 | **-0.15** | **0.027** | -0.05 | 0.442 | **-0.18** | **0.021** |

| | | | | | | | | | | | | |
|---|---|---|---|---|---|---|---|---|---|---|---|---|
| 16 | 10706328 | rs12149828 | Imputed | 0.97 | A/G | 0.17 | 0.08 | 0.314 | 0.07 | 0.316 | 0.02 | 0.809 |
| 16 | 58075282 | rs12447804 | Genotyped | 0.99 | T/C | 0.18 | 0.09 | 0.244 | 0.02 | 0.810 | 0.15 | 0.079 |
| 16 | 75390316 | rs2865531 | Imputed | 1.00 | A/T | 0.60 | -0.07 | 0.239 | -0.02 | 0.720 | -0.07 | 0.302 |
| 16 | 78187138 | rs1079572 | Genotyped | 1.00 | A/G | 0.63 | 0.03 | 0.616 | 0.04 | 0.510 | 0.00 | 0.943 |
| 17 | 68976415 | rs6501431 | Genotyped | 1.00 | T/C | 0.75 | 0.02 | 0.751 | -0.05 | 0.448 | 0.08 | 0.258 |
| 19 | 41124155 | rs113473882 | Imputed | 0.99 | C/T | 0.01 | -0.03 | 0.911 | -0.34 | 0.132 | 0.47 | 0.081 |
| 21 | 35652239 | rs9978142 | Imputed | 0.97 | T/A | 0.16 | 0.08 | 0.299 | 0.03 | 0.654 | 0.08 | 0.354 |
| 22 | 28056338 | rs134041 | Imputed | 0.99 | C/T | 0.58 | 0.02 | 0.727 | -0.02 | 0.668 | 0.06 | 0.351 |

All associations between genotype status at single SNPs (additive model) and change in lung function z-scores were adjusted for principal components (PCs) 1–10 and age as covariates in the linear mixed models with a random participant intercept. **Bold**: nominal significant association (p-value < 0.05)

Chr: chromosome; Pos: position; $R^2$: estimated value of the squared correlation between imputed genotypes and true, unobserved genotypes; EA/OA: effect allele / other allele; EA Freq: EA Frequency; NA: not available

**Table S7:** Association between baseline BMI and change in lung function z-scores within the study period (N=601).

| **Change in lung function** | **β-estimates (95%-CI)** | **p-value** |
|---|---|---|
| $FEV_1$ | 0.00 (-0.02; 0.02) | 0.982 |
| FVC | 0.00 (-0.02; 0.01) | 0.711 |
| $FEV_1$/FVC | 0.01 (-0.01; 0.02) | 0.534 |

β-estimates, 95% confidence intervals (95%-CI) and p-values per gaining 1 point in BMI. Adjusted for age, level of education, smoking (categorized as current, former or never smoking), exposure to second hand smoke (SHS) and change in $NO_2$ within the follow-up period. **Bold**: significant association (p-value < 0.05)

**Table S8:** Association between average BMI and change in lung function z-scores within the study period (N=601).

| Change in lung function | β-estimates (95%-CI) | p-value |
|---|---|---|
| $FEV_1$ | 0.00 (-0.02; 0.01) | 0.716 |
| FVC | -0.01 (-0.02; 0.00) | 0.181 |
| $FEV_1$/FVC | 0.01 (0.00; 0.03) | 0.151 |

β-estimates, 95% confidence intervals (95%-CI) and p-values per gaining 1 point in BMI. Adjusted for age, level of education, smoking (categorized as current, former or never smoking), exposure to second hand smoke (SHS) and change in $NO_2$ within the follow-up period. **Bold**: significant association (p-value < 0.05)

**Table S9:** Association between improvement of air quality and change in lung function z-scores stratified by baseline BMI (N=601).

| **Change in lung function** | **Improvement in exposure** | **1st tertile BMI (<23.8)** | **2nd tertile BMI (23.8 to 28.7)** | **3rd tertile BMI (>28.7)** | **p-value continuous interaction** | **p-value interaction (1st vs. 2nd)** | **p-value interaction (2nd vs. 3rd)** |
|---|---|---|---|---|---|---|---|
| $FEV_1$ | $NO_2$ (per 10 µg/m$^3$) | 0.09 (-0.08; 0.26) | **0.22 (0.05; 0.39)** | 0.07 (-0.10; 0.25) | 0.974 | 0.208 | 0.174 |
| | $NO_x$ (per 20 µg/m$^3$) | 0.04 (-0.08; 0.16) | **0.15 (0.03; 0.27)** | 0.02 (-0.11; 0.15) | 0.867 | 0.182 | 0.142 |
| | $PM_{2.5}$ (per 5 µg/m$^3$) | -0.07 (-0.19; 0.05) | 0.05 (-0.07; 0.17) | -0.07 (-0.21; 0.07) | 0.819 | 0.085 | 0.127 |
| | $PM_{10}$ (per 10 µg/m$^3$) | 0.05 (-0.12; 0.22) | **0.19 (0.02; 0.36)** | 0.05 (-0.15; 0.25) | 0.970 | 0.174 | 0.204 |
| FVC | $NO_2$ (per 10 µg/m$^3$) | -0.01 (-0.17; 0.15) | 0.05 (-0.11; 0.21) | -0.06 (-0.23; 0.11) | 0.656 | 0.574 | 0.323 |
| | $NO_x$ (per 20 µg/m$^3$) | 0.01 (-0.10; 0.12) | 0.04 (-0.07; 0.16) | -0.04 (-0.17; 0.09) | 0.554 | 0.692 | 0.322 |
| | $PM_{2.5}$ (per 5 µg/m$^3$) | -0.03 (-0.14; 0.08) | 0.02 (-0.09; 0.14) | -0.12 (-0.25; 0.01) | 0.226 | 0.44 | 0.058 |
| | $PM_{10}$ (per 10 µg/m$^3$) | -0.05 (-0.21; 0.11) | 0.03 (-0.13; 0.19) | -0.16 (-0.35; 0.03) | 0.208 | 0.415 | 0.075 |
| $FEV_1$/FVC | $NO_2$ (per 10 µg/m$^3$) | 0.14 (-0.05; 0.33) | **0.24 (0.06; 0.43)** | **0.21 (0.02; 0.41)** | 0.468 | 0.359 | 0.815 |
| | $NO_x$ (per 20 µg/m$^3$) | 0.04 (-0.09; 0.17) | **0.14 (0.01; 0.28)** | 0.09 (-0.06; 0.24) | 0.556 | 0.229 | 0.556 |
| | $PM_{2.5}$ (per 5 µg/m$^3$) | -0.06 (-0.19; 0.08) | 0.01 (-0.12; 0.14) | 0.08 (-0.07; 0.24) | 0.108 | 0.381 | 0.434 |
| | $PM_{10}$ (per 10 µg/m$^3$) | 0.16 (-0.03; 0.35) | **0.21 (0.02; 0.40)** | **0.34 (0.11; 0.56)** | 0.120 | 0.675 | 0.306 |

β-estimates and 95% confidence intervals (95%-CI) per an improvement of 10 µg/m$^3$ in $NO_2$, 20 µg/m$^3$ in $NO_x$, 5 µg/m$^3$ in $PM_{2.5}$ and 10 µg/m$^3$ in $PM_{10}$. P-values are given for the interaction terms between the continuous BMI measurements and air pollution (p-value continuous interaction) as well as for the interaction terms between tertiles of average BMI and air pollution. Adjusted for age, level of education, smoking (categorized as current, former or never smoking) and exposure to second hand smoke (SHS). **Bold**: significant association (p-value < 0.05)

### References


1. Eeftens M, Beelen R, de Hoogh K, Bellander T, Cesaroni G, Cirach M, Declercq C, Dėdelė A, Dons E, de Nazelle A, Dimakopoulou K, Eriksen K, Falq G, Fischer P, Galassi C, Gražulevičienė R, Heinrich J, Hoffmann B, Jerrett M, Keidel D, Korek M, Lanki T, Lindley S, Madsen C, Mölter A, Nádor G, Nieuwenhuijsen M, Nonnemacher M, Pedeli X, Raaschou-Nielsen O, et al. Development of Land Use Regression Models for PM 2.5 , PM 2.5 Absorbance, PM 10 and PM coarse in 20 European Study Areas; Results of the ESCAPE Project. *Environ. Sci. Technol.* American Chemical Society; 2012; 46: 11195–11205.

2. Beelen R, Hoek G, Vienneau D, Eeftens M, Dimakopoulou K, Pedeli X, Tsai M-Y, Künzli N, Schikowski T, Marcon A, Eriksen KT, Raaschou-Nielsen O, Stephanou E, Patelarou E, Lanki T, Yli-Tuomi T, Declercq C, Falq G, Stempfelet M, Birk M, Cyrys J, von Klot S, Nádor G, Varró MJ, Dėdelė A, Gražulevičienė R, Mölter A, Lindley S, Madsen C, Cesaroni G, et al. Development of NO2 and NOx land use regression models for estimating air pollution exposure in 36 study areas in Europe – The ESCAPE project. *Atmos. Environ.* 2013; 72: 10–23.

3. Das S, Forer L, Schönherr S, Sidore C, Locke AE, Kwong A, Vrieze SI, Chew EY, Levy S, McGue M, Schlessinger D, Stambolian D, Loh PR, Iacono WG, Swaroop A, Scott LJ, Cucca F, Kronenberg F, Boehnke M, Abecasis GR, Fuchsberger C. Next-generation genotype imputation service and methods. *Nat. Genet.* 2016; 48: 1284–1287.

4. Soler Artigas M, Wain L V, Miller S, Kheirallah AK, Huffman JE, Ntalla I, Shrine N, Obeidat M, Trochet H, McArdle WL, Alves AC, Hui J, Zhao JH, Joshi PK, Teumer A, Albrecht E, Imboden M, Rawal R, Lopez LM, Marten J, Enroth S, Surakka I, Polasek O, Lyytikäinen L-P, Granell R, Hysi PG, Flexeder C, Mahajan A, Beilby J, Bossé Y, et al. Sixteen new lung function signals identified through 1000 Genomes Project reference panel imputation. *Nat. Commun.* 2015; 6: 8658.
